## Supplementary figures and images for "Single-cell transcriptomic profiling of *C. elegans* Q neuroblast lineage during migration and differentiation"

### S1 Fig

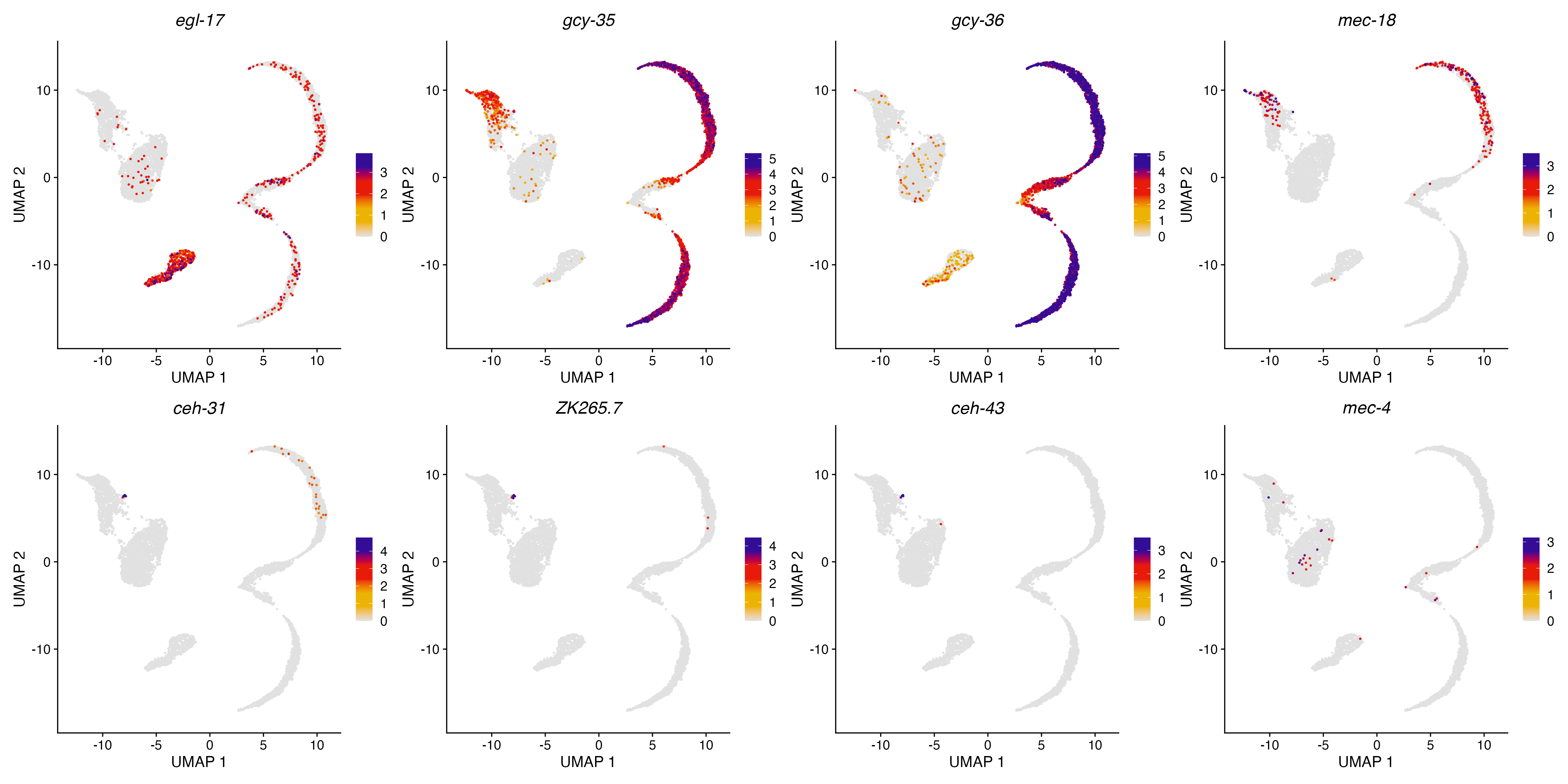

### S2 Fig

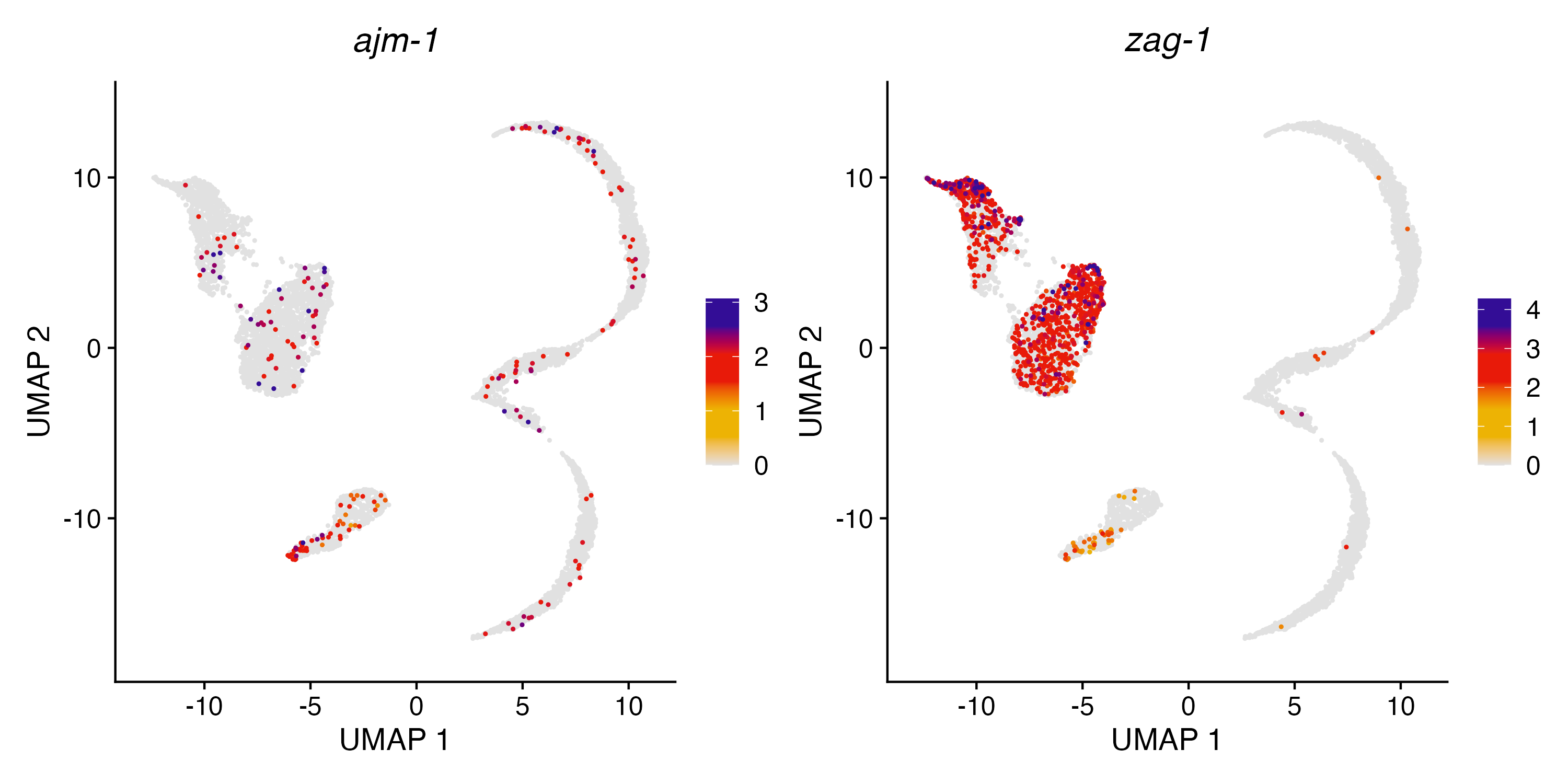

### S3 Fig Fig

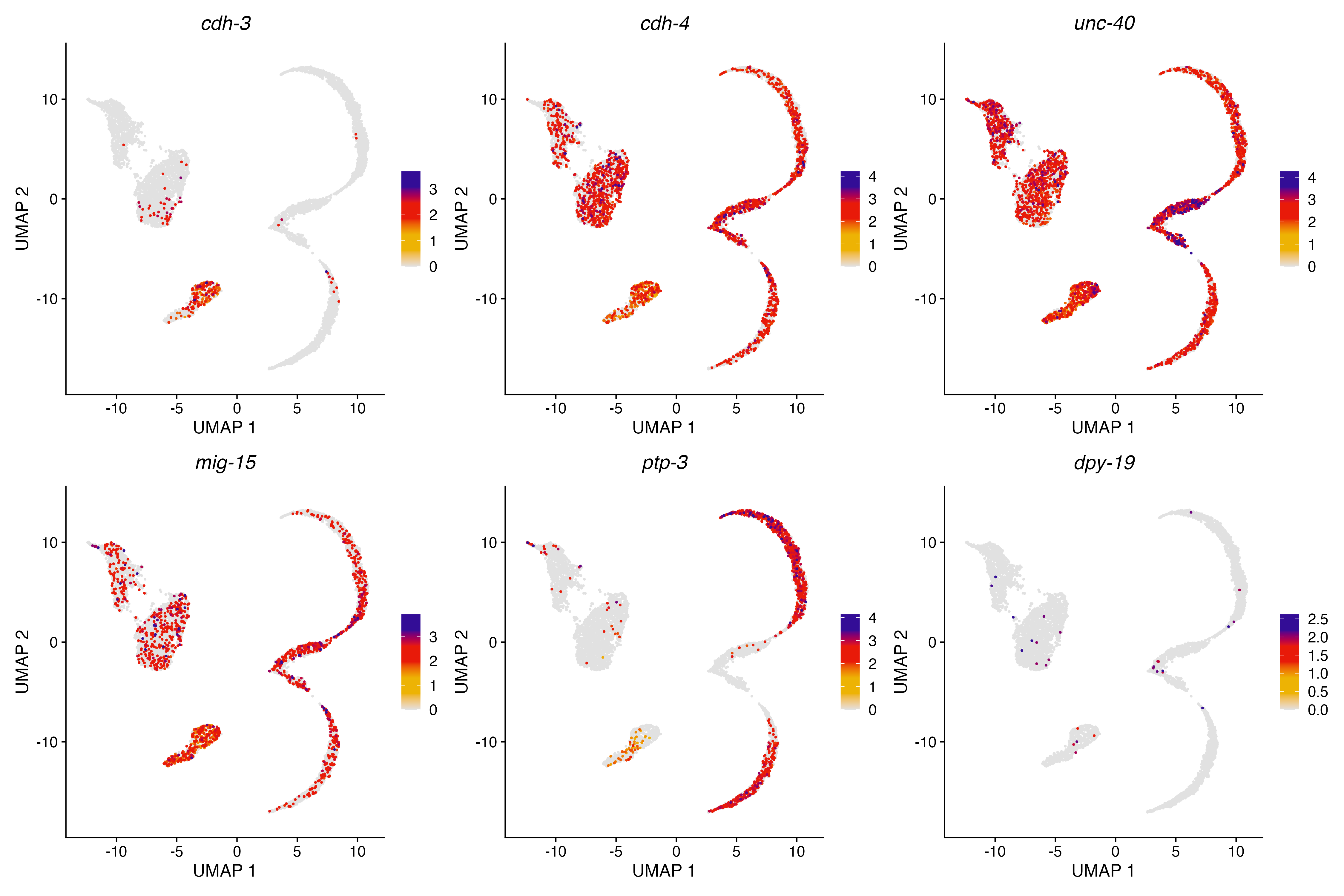

### S4 Fig Fig

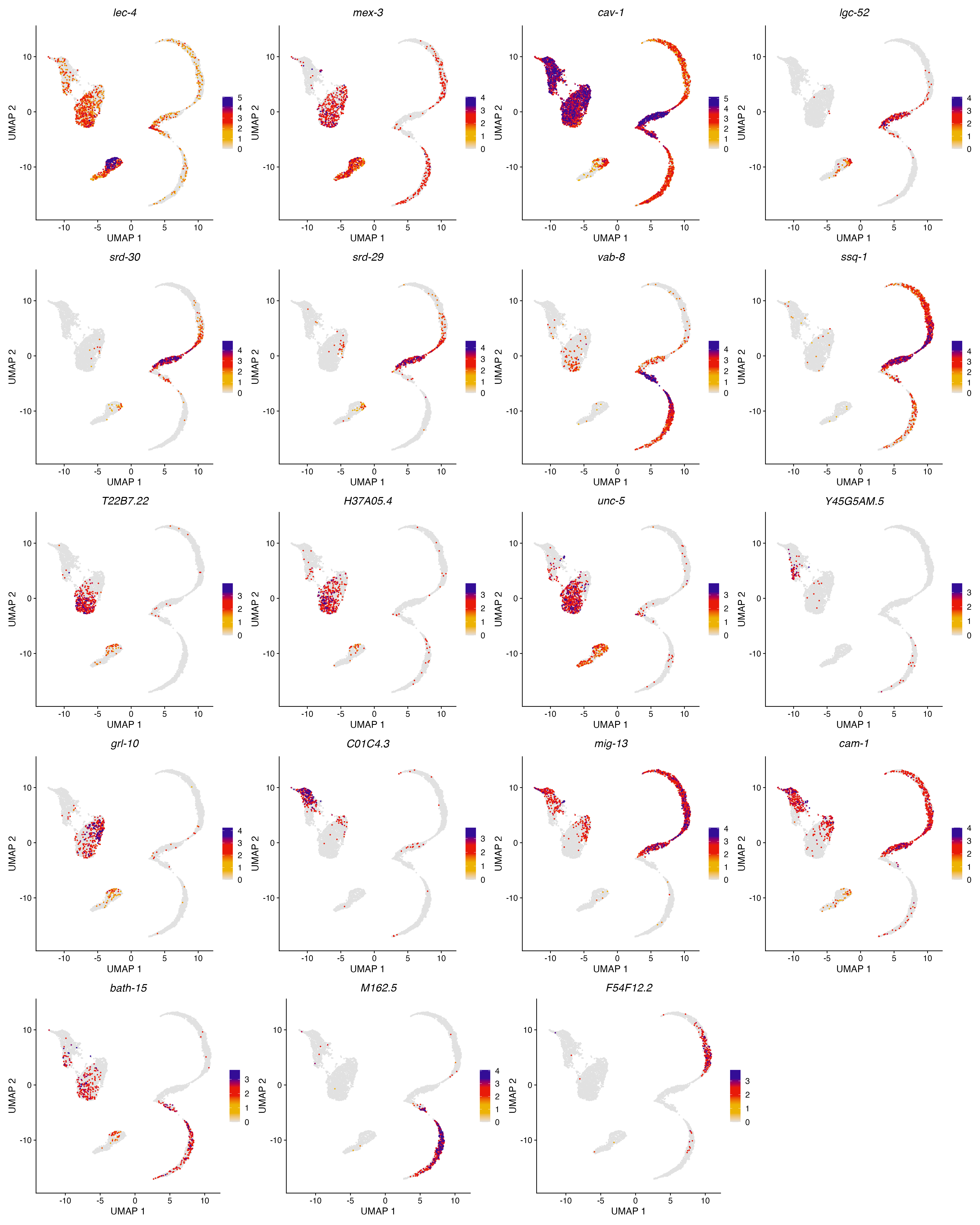

### S5 Fig

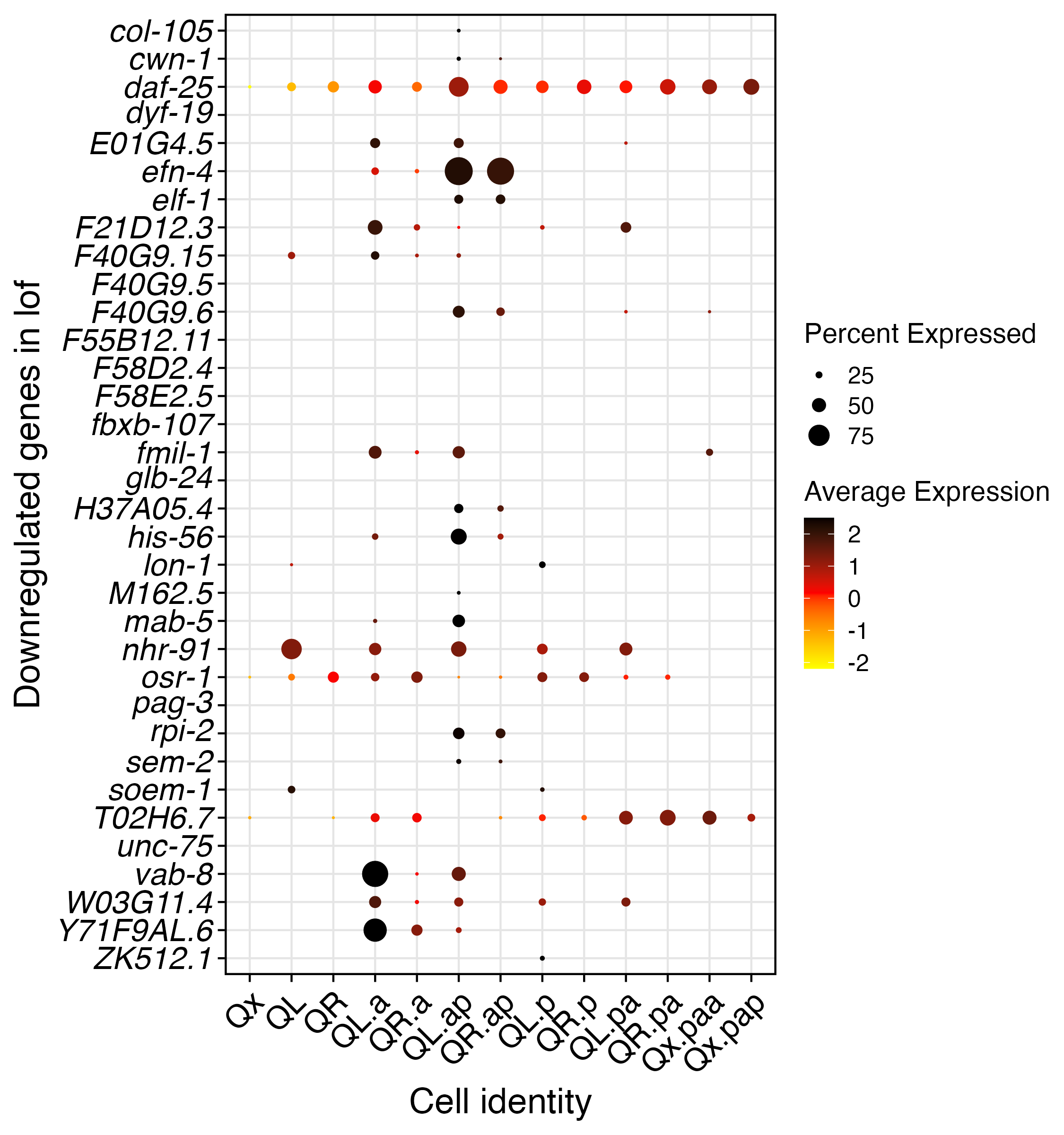

### S6 Fig

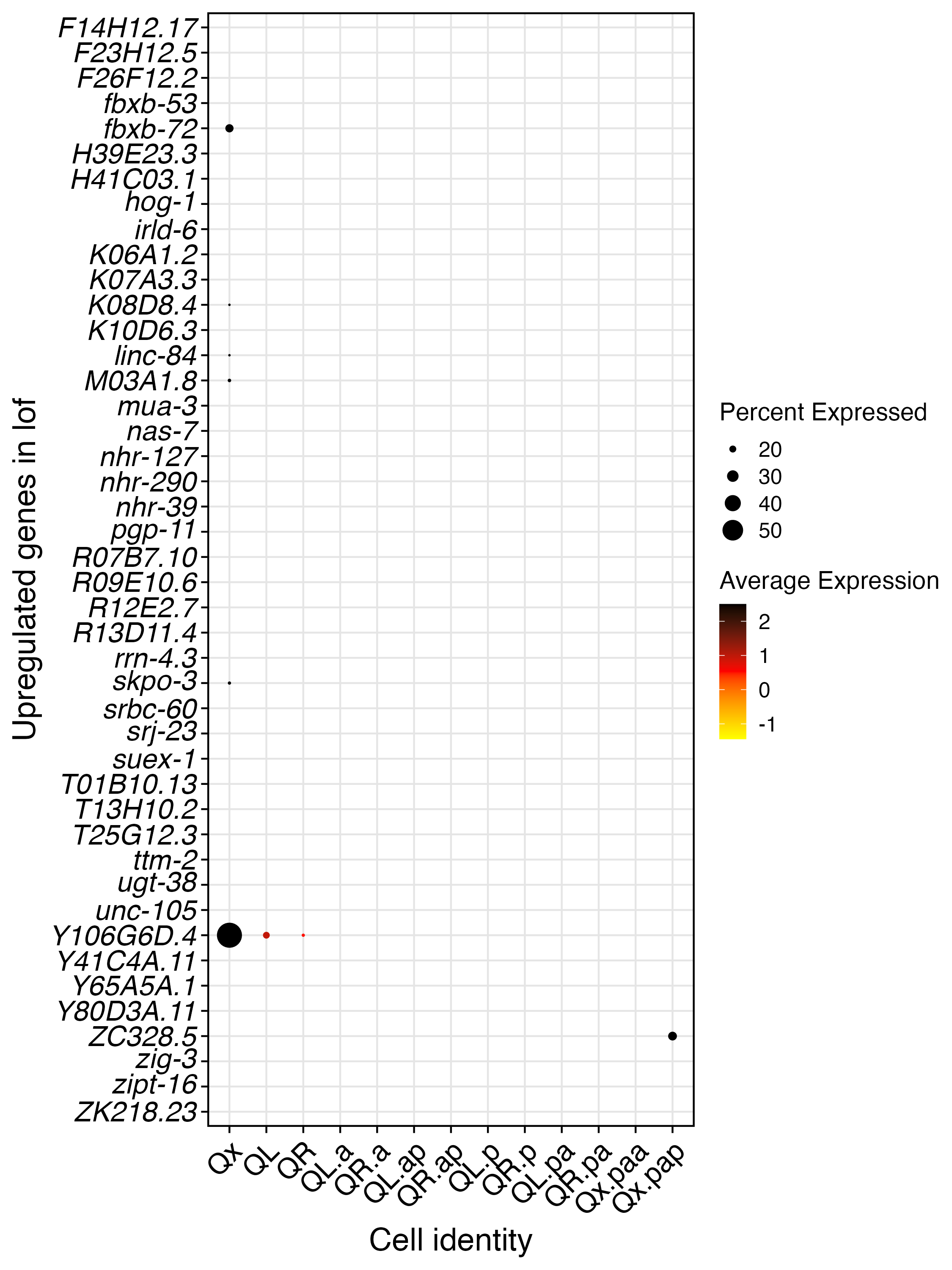

### S7 Fig

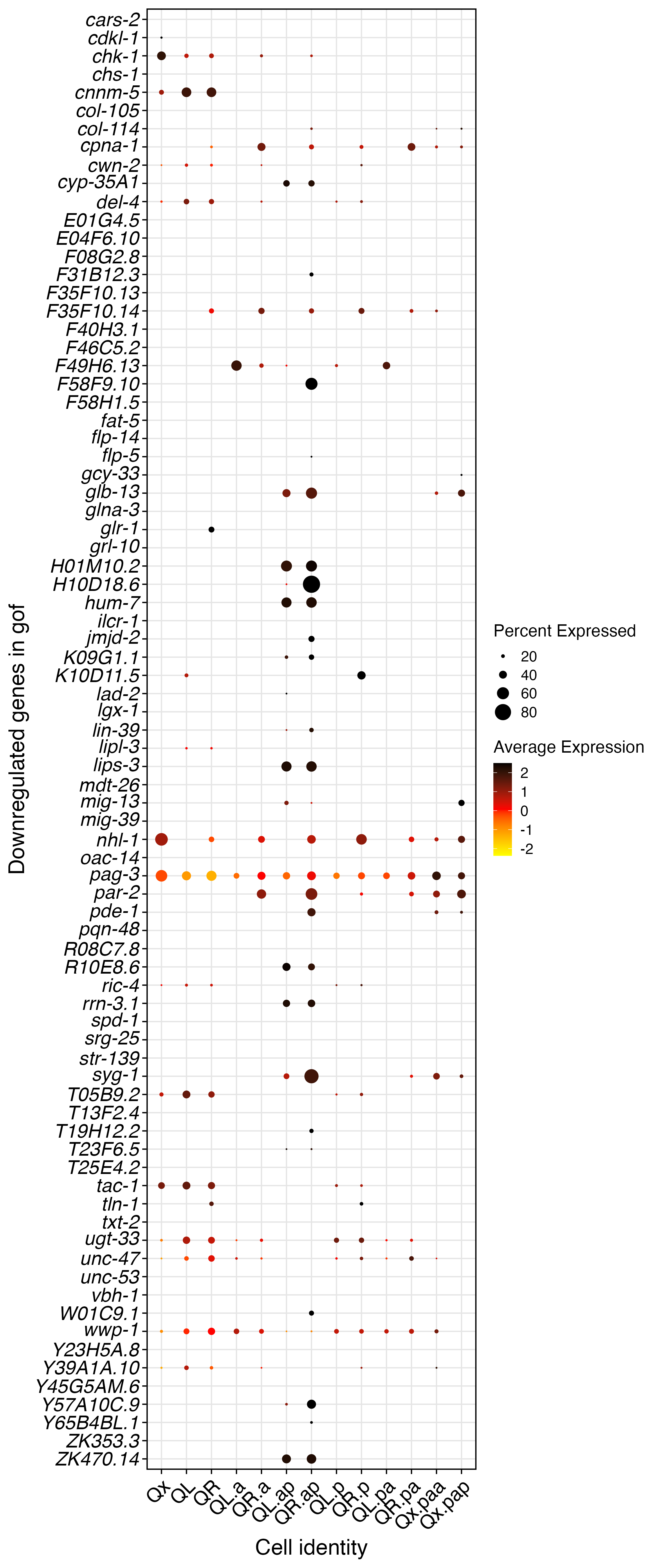

### S8 Fig

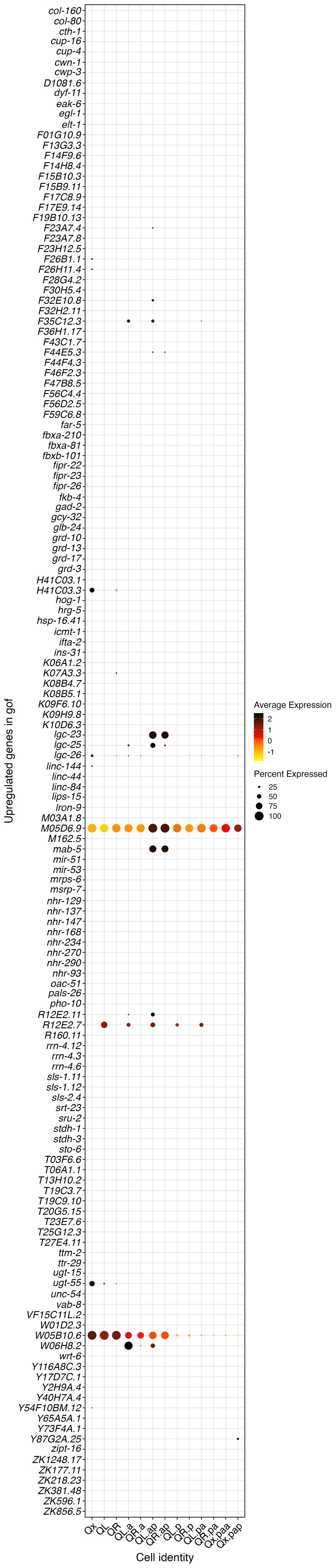

### S9 Fig

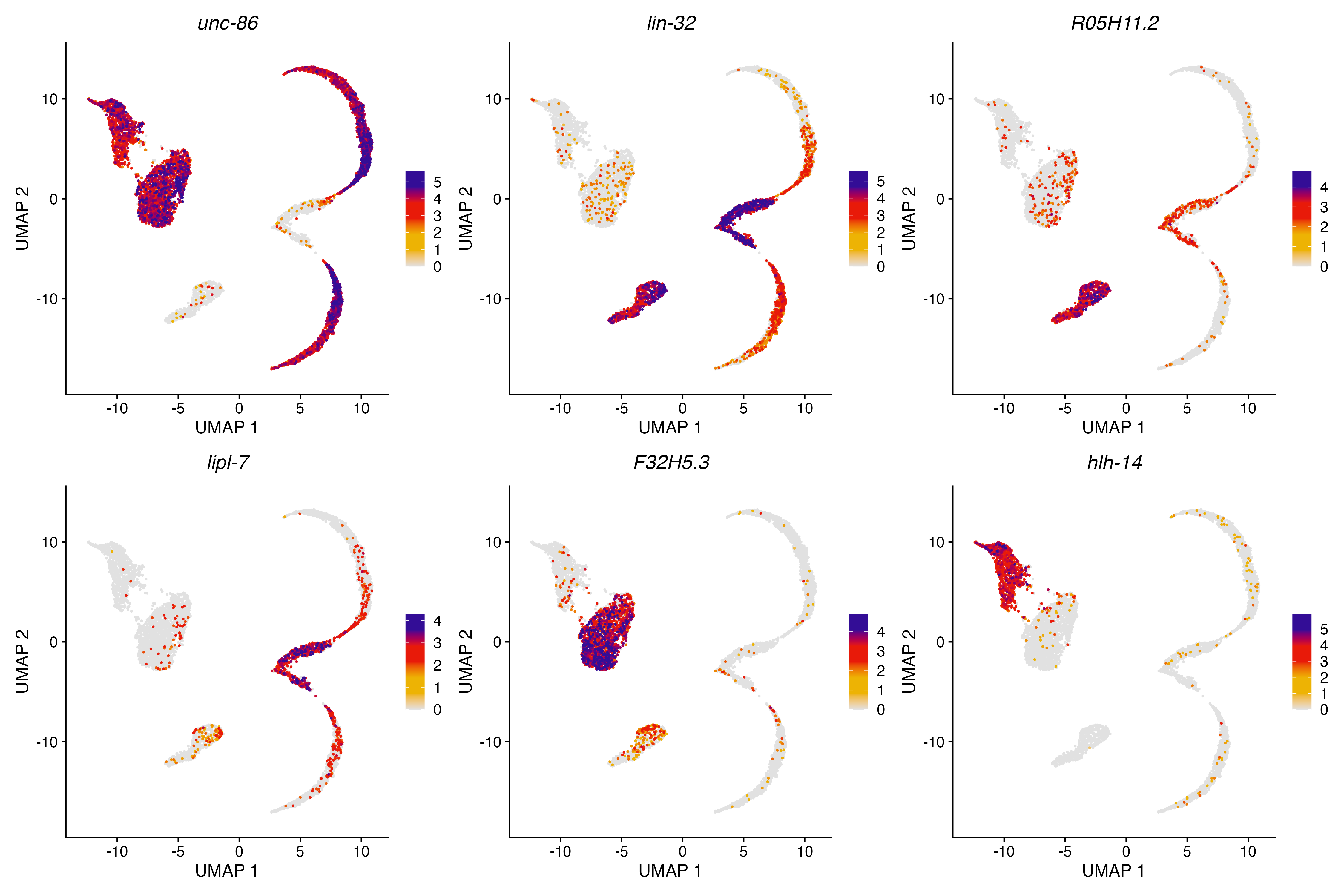

### S10 Fig

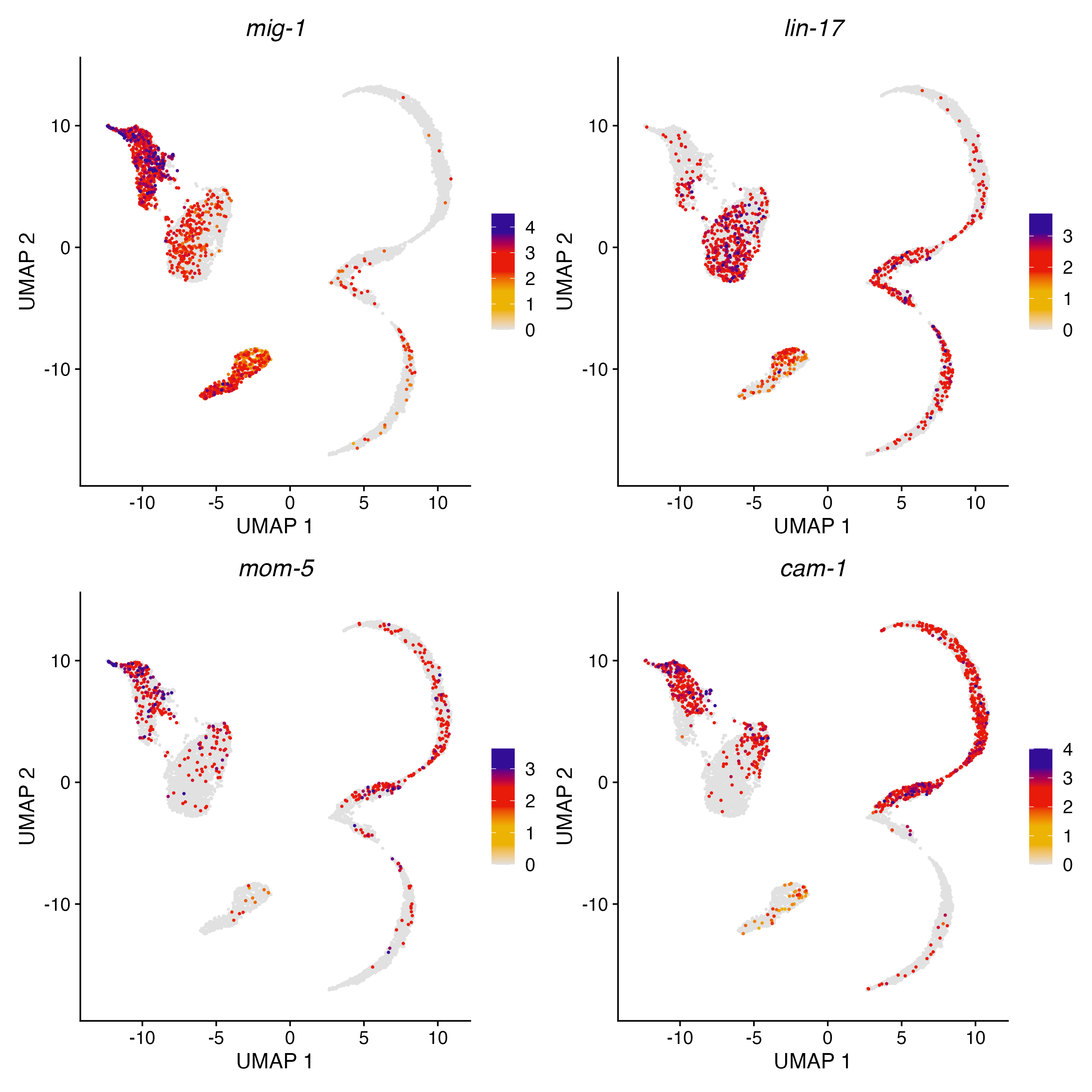
